# Genome sequence of *Pseudopithomyces chartarum*, causal agent of facial eczema (pithomycotoxicosis) in ruminants, and identification of the putative sporidesmin toxin gene cluster

**DOI:** 10.1101/2021.04.29.441555

**Authors:** Jaspreet Singh Sidhu, Vinod Suresh, Abdul Baten, Ann M. McCartney, Gavin Lear, Jan M. Sprosen, Mark H. Oliver, Natasha T. Forester, Paul H. Maclean, Nikola Palevich, Ruy Jauregui, Christine R. Voisey

**Affiliations:** AgResearch Limited, Grasslands Research Centre, Palmerston North, New Zealand; Department of Engineering Science, University of Auckland, Auckland, New Zealand; Auckland Bioengineering Institute, University of Auckland, Auckland, New Zealand; Manaaki Whenua – Landcare Research, Auckland, New Zealand; School of Biological Sciences, University of Auckland, Auckland, New Zealand; AgResearch Limited, Ruakura Research Centre, Hamilton, New Zealand; Liggins Institute, University of Auckland, New Zealand; Institute of Precision Medicine & Bioinformatics, Sydney Local Health District, Royal Prince Alfred Hospital, Camperdown, NSW 2050, Australia

**Keywords:** Epipolythiodioxopiperazine, non-ribosomal peptide synthetase, flavin-dependent halogenase, *Pseudopithomyces chartarum*, mycotoxin, animal health

## Abstract

Facial eczema (FE) in grazing ruminants is a debilitating liver syndrome induced by ingestion of sporidesmin, a toxin belonging to the epipolythiodioxopiperazine class of compounds. Sporidesmin is produced in spores of the fungus *Pseudopithomyces chartarum*, a microbe which colonises leaf litter in pastures. New Zealand has a high occurrence of FE in comparison to other countries as animals are fed predominantly on ryegrass, a species that supports high levels of *Pse. chartarum* spores. The climate is also particularly conducive for *Pse. chartarum* growth. Here, we present the genome of *Pse. chartarum* and identify the putative sporidesmin gene cluster. The *Pse. chartarum* genome was sequenced using single molecule real-time sequencing (PacBio) and gene models identified. Loci containing genes with homology to the aspirochlorine, sirodesmin PL and gliotoxin cluster genes of *Aspergillus oryzae, Leptosphaeria maculans* and *Aspergillus fumigatus*, respectively, were identified by tBLASTn. We identified and annotated an epipolythiodioxopiperazine cluster at a single locus with all the functionality required to synthesise sporidesmin.

**Highlights:**

- The whole genome of *Pseudopithomyces chartarum* has been sequenced and assembled.
- The genome is 39.13 Mb, 99% complete, and contains 11,711 protein coding genes.
- A putative sporidesmin A toxin (cause of facial eczema) gene cluster is described.
- The genomes of *Pse. chartarum* and the *Leptosphaerulina chartarum* teleomorph differ.
- Comparative genomics is required to further resolve the *Pseudopithomyces* clade.

## 1. Introduction

Pithomycotoxicosis (also known as facial eczema [FE]) is a serious liver syndrome in ruminants grazing ryegrass pastures containing spores of the dothideomycete fungus *Pseudopithomyces chartarum sensu lato* (Berk. & M.A. Curtis) Jun F. Li, Ariyaw. & K.D. Hyde, formerly *Pithomyces chartarum* (Berk. & M.A. Curtis) [1-3]. The spores contain sporidesmin (Fig. 1), an epipolythiodioxopiperazine (ETP) toxin, which causes liver toxicity when ingested by ruminants. Sporidesmin is a collective term encompassing nine related compounds (A-H and J), with sporidesmin A the most toxic and constituting 90% of the sporidesmins produced in culture [4]. Sporidesmin levels are not always correlated with spore production and can vary according to culture conditions [11]. Autooxidation of sporidesmin A in the liver induces severe local free radical-induced damage and accumulation of photoreactive chlorophyll degradation byproducts which induce FE. Cholestatic liver damage resulting from consumption of spore-contaminated feed results in impaired reproductive indices and animal productivity [12] well before facial lesions are seen at near end-state pathology [3].

**Fig. 1.**
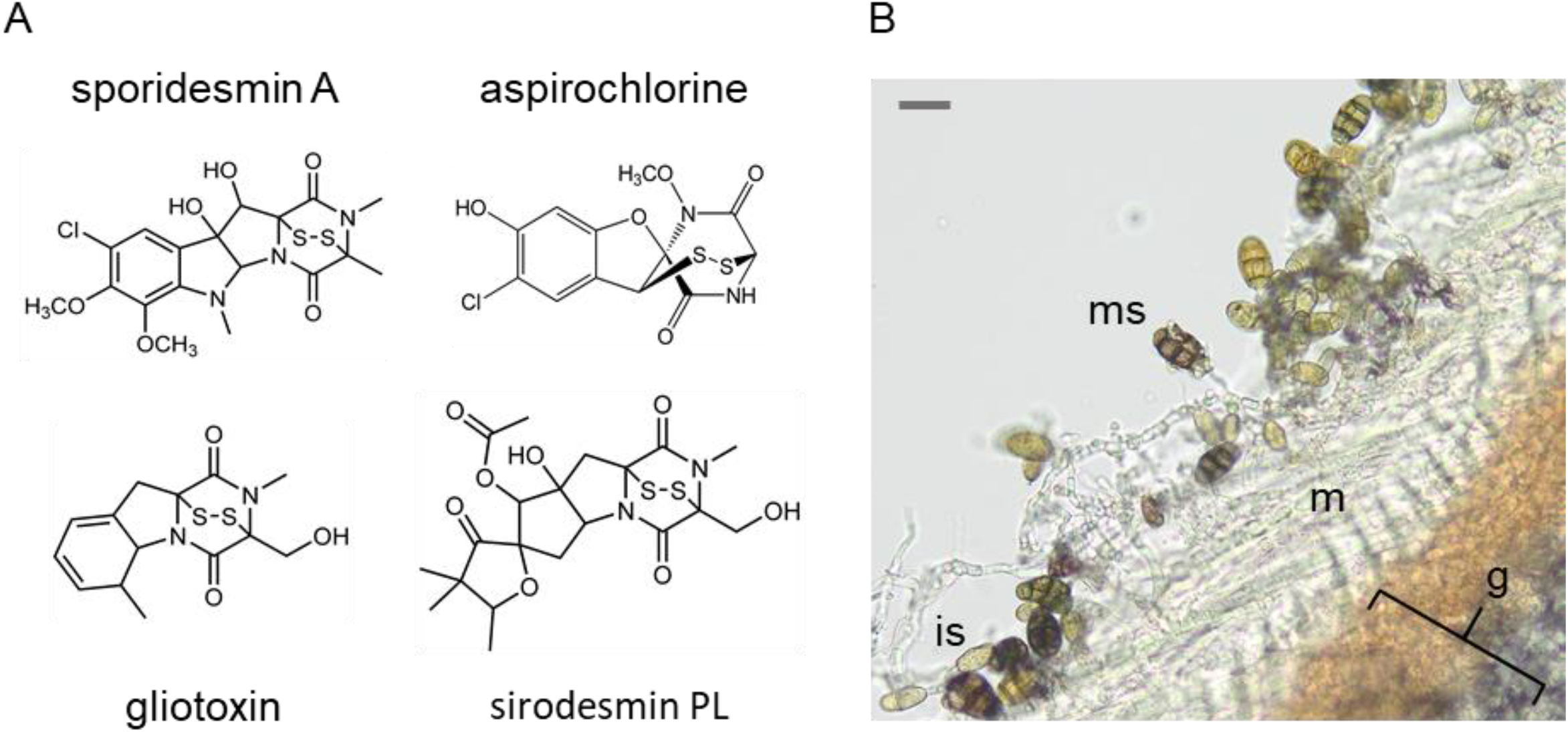
Schematic diagrams of sporidesmin A and other ETP toxins, and *Pse. chartarum* 91/35/29 spore development. (A) Structures of sporidesmin A (*Pse. chartarum*), aspirochlorine (*A. oryzae*), sirodesmin PL (*L. maculans*) and gliotoxin (*A. fumigatus*) showing the characteristic ETP disulphide bridge and the chlorinated moiety (sporidesmin A and aspirochlorine only). (B) Bright field micrograph of a transverse section through a wheat grain (g) infected with *Pse. chartarum*. Mycelia (m) after 3 weeks of growth at 22 °C are shown. Immature spores (is) are small and pale and lack obvious septa, and mature spores (ms) have heavily melanised longitudinal and transverse septa. The section is 30 µm thick and the scale bar is 20 µm.

*Pse. chartarum* has been reported as both an endophyte and plant pathogen in many regions of the world [6-9]; it is also a saprophyte on decaying leaf litter, including pasture grasses such as perennial ryegrass (*Lolium perenne* L.) [2, 10]. Non-toxigenic strains are common and their proportion varies widely between and within countries [9]. In countries with a high proportion of toxigenic strains, and a temperate climate, the FE burden on grass-fed animals is high and is estimated to reduce production losses in affected countries significantly (e.g. >NZD100M per annum in New Zealand) [5]. Current FE control methods such as fungicide application to pastures [13], and very high animal dosing with zinc [14] have some efficacy but disrupt copper and molybdenum regulation in ruminants [15] and have adverse consequences for soils and waterways [2].

ETPs are a class of secondary metabolites, made only by fungi, that are characterised by the presence of a sulphur-bridged dioxopiperazine ring synthesised from two amino acids by a non-ribosomal peptide synthetase (NRPS). ETP toxicity is attributed to the generation of reactive oxygen species (ROS) through redox cycling and cross-linking to proteins through their cysteine residues which causes cellular damage [3, 16]. Putative ETP gene clusters have been found in many ascomycete taxa [17] and include well-characterised examples such as the sirodesmin (*sir*) [18], gliotoxin (*gli*) [19], and aspirochlorine (*acl*) [20] clusters of *L. maculans, A. fumigatus*, and *A. oryzae* respectively. Similar to sporidesmin, aspirochlorine is halogenated and the *acl* cluster contains a flavin-dependent halogenase (*aclH*) which chlorinates the molecule at the last step of its synthesis [20].

Here we report the first genome sequence of a sporidesmin-producing strain of *Pse. chartarum*, originally isolated from ryegrass-based pasture in the North Island of New Zealand. ETP genes from the functionally characterised ETP clusters producing aspirochlorine, sirodesmin PL and gliotoxin were used to probe the assembled genome for potential ETP clusters. We have identified and described a single locus with all the genes required to synthesise a chlorinated ETP such as sporidesmin.

## 2. Results and Discussion

### 2.1. Genomic features of *Pse. chartarum* 91/35/29

Alignment of the ribosomal long subunit (LSU) gene from *Pse. chartarum* 91/35/29 with the LSU of other members of the *Didymosphaeriaceae* (Pleosporales) [21] places *Pse. chartarum* 91/35/29 in the same clade as *Pse. chartarum* C459 (an isolate from fruit), as well as *L. chartarum (the teleomorph of Pse. chartarum)*, an isolate from *Nicotiana tabacum* (Fig. S1). The genome of *Pse. chartarum* 91/35/29, assembled exclusively from 20 kb (SMRT [Sequel]) PacBio reads, has an estimated nuclear genome size of 39.13 Mbp and a mitochondrial genome of 101.5 Kbp, with an overall G + C content of 50.16% (Table 1). The resolved genome consists of 22 scaffolds (N50 = 2.3 Mbp) with high coverage, approximately 140-fold. The *Pse. chartarum* genome is similar in size to the median assembly size (36.5) Mbp) of 101 *Dothideomycetes* genomes published recently through the 1000 Fungal Genomes Project, although genome size in the *Dothideomycetes* can vary widely (between 16.95 Mbp and 177.60 Mbp) [22].

**Table 1.** Comparison statistics for the *Pse. chartarum* 91/35/29 and *L. chartarum* genomes.

| Feature | *Pse. chartarum* | *L. chartarum* |
| --- | --- | --- |
| **Genome project information** |  |  |
| Status | High-quality draft | Draft |
| Isolation source | Ryegrass | Tobacco |
| BioSample ID | SUB6703657 | N/A |
| BioProject ID | PRJNA596299 | PRJNA400396 |
| Assembly method | CANU | SPAdes |
| Genome coverage | 140× | N/A |
| Sequencing technology | PacBio | Illumina |
| **Scaffold statistics** |  |  |
| Assembled genome size (Mb) | 39.1 | 37.9 |
| Scaffold count | 22 | 18,713 |
| Longest scaffold (Mb) | 3.6 | 1.4 |
| Scaffold N50 (Mb) | 2.3 | 0.3 |
| Number of N50 contigs | 7 | 35 |
| Scaffold N90 (bp) | 1.5 | 0.1 |
| Number of N90 contigs | 16 | 333 |
| **Assembly base composition** |  |  |
| DNA G+C content (%) | 50.2 | 59.4 |
| DNA A+T content (%) | 49.9 | 40.6 |
| DNA N (%) | 0.00 | 0.01 |
| **Gene model and genome assembly completeness** |  |  |
| Protein coding genes | 11,711 | 15,091 |
| BUSCO: C:P:M (%)* | 99:0.0:1.0 | 98.9:0.3:0.8 |
| Reference | This report | [6] |

\*C, P, and M refer to fully represented, partially represented, and missing BUSCO genes [35]. N/A, used where data is not available.

The nearest sequenced genome is from a proposed *Pse. chartarum* teleomorph, *L chartarum*, sequenced on the Illumina HiSeq platform [6]. As expected, *L. chartarum* has a similar genome size to *Pse. chartarum* (37.9 Mbp; Table 1), however the G + C content of *L. chartarum* was substantially higher than *Pse. Chartarum* (59.4% versus 50.19%). *Pse. chartarum* strains have been isolated from a range of monocotyledonous and dicotyledonous species [23-25] and have also occasionally been recovered from human clinical specimens [26]. They have variously been recorded as having saprophytic, endophytic or pathogenic lifestyles [24]. Comparative genomics is therefore required to resolve the taxonomy of this group as, to our knowledge, the *Pse. chartarum* and *L. chartarum* isolates are the only representatives with a sequenced genome [23-26].

*Ab initio* gene prediction resulted in 11,711 annotated protein coding regions (PCGs) in *Pse. chartarum*, which is consistent for a fungal genome of this size [27] and similar to the average gene prediction (12,720) from the 101 *Dothideomycetes* genomes referred to above [22]. The number of PCGs for *L. chartarum* was predicted to be 15,091, further supporting the likelihood that these fungi are genetically quite distinct. Benchmarking Universal Single-Copy Orthologs (BUSCO) pipelines revealed that *Pse. chartarum* 91/35/29 genome assembly and annotation was 99.0% complete when searched against the fungal reference database (Table 1).

### 2.2. Genomic insight into the putative sporidesmin gene cluster

A key aim of sequencing *Pse. chartarum* 91/35/29 was to identify the putative sporidesmin gene cluster. The first step was to align the aspirochlorine (*acl*), sirodesmin (*sir*) and gliotoxin (*gli*) cluster genes against the *Pse. chartarum* genome to identify loci containing putative homologs of ETP genes (Tables S1A, B and C). The three comparisons revealed a single *Pse. chartarum* genomic locus that contained all the genes required to make a chlorinated ETP. The cluster is approximately 43.3 kb in length and contains 21 genes (*spd1*- *spd21*) (Table S2). Next, the corresponding protein sequences were extracted, their predicted domain architecture determined by Interpro and Pfam, and BLASTP used to find related sequences in the Uniprot/Swissprot database. Twelve of the inferred proteins in the cluster had highest homology to enzymes involved in aspirochlorine (AclH, AclG, AclQ, AclK, AclN, AclD, AclI, AclA and AclM), sirodesmin PL (SirJ and SirB) and gliotoxin (GliP) synthesis, although amino acid identity was generally low (29.6.%- 60.3%; Table S2). Spd proteins with the same domain architecture as *acl, sir* or *gli* cluster proteins were deemed to be likely orthologues or functional equivalents.

The sporidesmin cluster (Fig. 2) contains a bimodular NRPS (Spd17; similar to AclP, SirP and GliP) predicted to condense two amino acids to form the sporidesmin core, plus a putative cytochrome P450 (Spd3; a likely orthologue of GliC), a glutathione-S-transferase (Spd5; a likely orthologue of AclG, SirG and GliG), a gamma-glutamyl cyclotransferase (Spd9; a likely orthologue of AclK and GliK), a dipeptidase (Spd2, a likely orthologue of AclJ, SirJ and GliJ), an amino transferase (Spd16; a likely orthologue of AclI, SirI and GliI) an d a thioredoxin reductase (Spd13; a likely orthologue of AclT, SirT and GliT), which collectively synthesise the sulphur bridge in ETPs [28-31]. Sporidesmin has a chlorine residue (Fig. 1A), and the presence of a putative flavin dependent halogenase (Spd 4), a likely orthologue of AclH which chlorinates aspirochlorine [20], strengthens the evidence that the cluster contains the genes for sporidesmin synthesis. Spd15 belongs to the acyl-CoA-acyltransferase superfamily (IPR016181), and a gene with this putative function is also in the sirodesmin (SirH) cluster, although the proteins are not conserved in amino acid identity or domain architecture. Similar to biosynthetic gene clusters (BGCs) in general, the putative sporidesmin cluster encodes several tailoring enzymes including methyl transferases (Spd1, Spd7, Spd11 and Spd21) and further cytochrome P450 monooxygenases (Spd8 and Spd10). Spd12 is a hypothetical protein with no known functional domains but has highest amino acid identity (60.3%) with the aspirochlorine biosynthesis protein AclN, a protein of the hydroxylase/desaturase family. The transporter-encoding genes in the Spd cluster include an ATP Binding Cassette-transporter (Spd6, a likely orthologue of AclQ and SirA, but not present in the gliotoxin cluster) and two Major Facilitator Superfamily (MFS) efflux transporters (Spd14 and Spd18), one of which (Spd18) has a similar domain architecture to AclA and GliA of *A. oryzae* and *A. fumigatus* respectively. Spd20 contains a cysteine-rich DNA binding motif and may regulate gene transcription. There are two hypothetical proteins in the cluster, Spd12 and Spd19, both with low scores to described proteins, and with no identified functional domains.

**Fig. 2.**
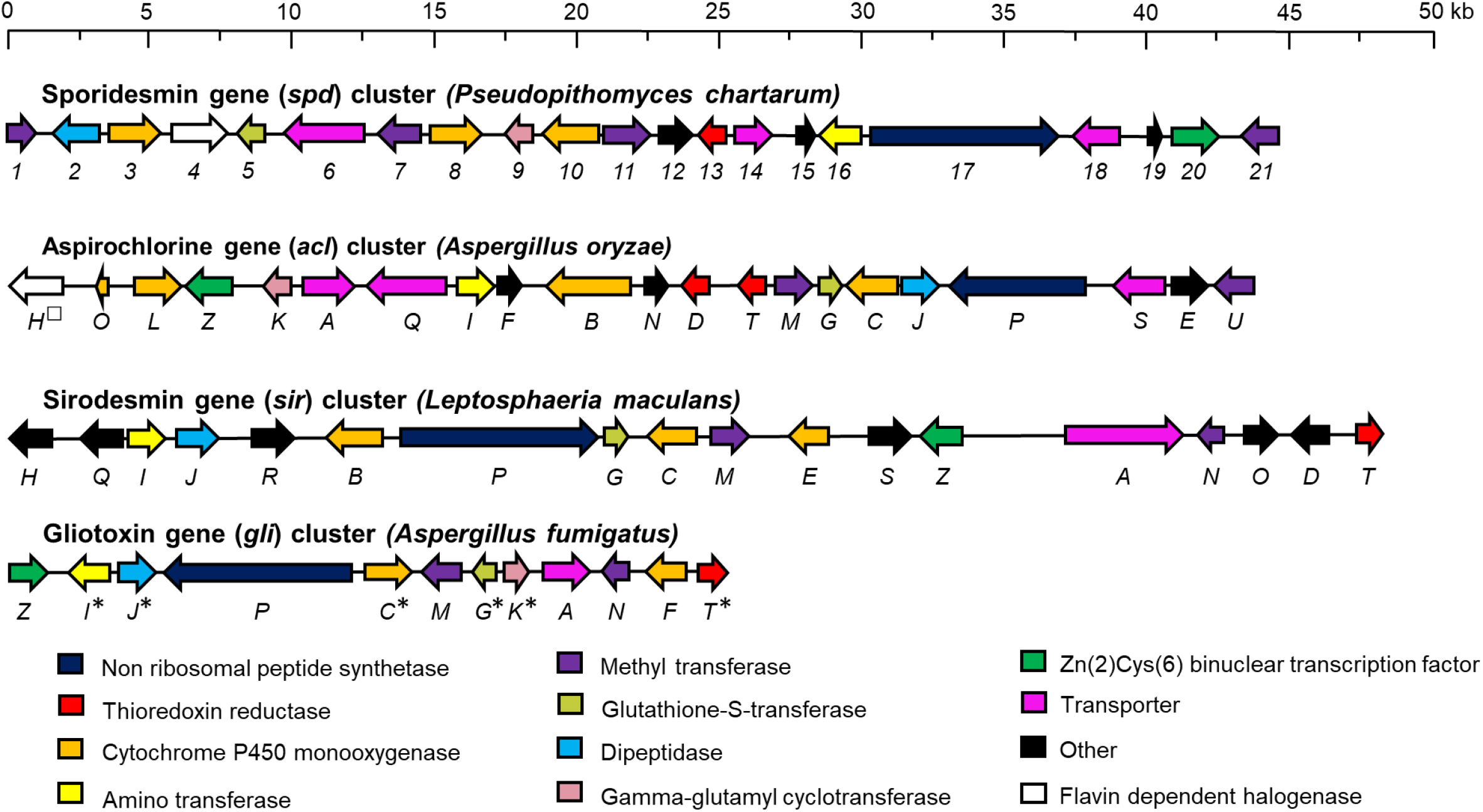
Cluster size and gene order of the putative sporidesmin (*spd*), aspirochlorine (*acl*), sirodesmin (*sir*) and gliotoxin (*gli*) BGCs. Clusters were left-justified, and features were drawn t o scale (see black bar above). The colour and orientation of arrows indicates proposed gene function and orientation respectively. Functionally characterised genes involved in gliotoxin disulphide bridge synthesis (asterisk) and addition of the chlorine residues (square) of aspirochlorine are marked.

## 3. Conclusions

We sequenced the genome of a sporidesmin-producing *Pse. chartarum* strain. Interrogation of the sequence with homologues of the *acl, sir* and *gli* genes points to this cluster being the only candidate with the functionality to synthesise sporidesmin. Functional characterisation of pathway genes is required to confirm that this cluster is responsible for sporidesmin synthesis. This discovery will be invaluable for determining the order of the chemical modifications through which sporidesmins A-J & I are synthesised, and may suggest strategies to disrupt accumulation of the most toxic analogue, sporidesmin A. Detailed sequence analysis is required to resolve the evolutionary relationships between *Pse. chartarum* isolates in the pasture environment, and knowledge of the gene cluster sequence will assist in determining why some *Pse. chartarum* strains do not synthesise sporidesmin, and permit the tracking of sporidesmin-producing strains in the environment.

## 4. Materials and methods

### 4.1. Pse. chartarum strain and storage conditions

The *Pse. chartarum* strain (91/35/29) sequenced in this study (kindly supplied by John Kerby, AgResearch Ltd., Ruakura Research Centre, Hamilton, New Zealand (NZ)) was isolated from field-grown ryegrass tissues at the Ruakura Research Centre. To grow the strain for long-term storage, a colony was incubated on potato dextrose agar (Difco™ Becton, Dickinson and Co. USA) in the dark at 22°C for 3 weeks. A 1 cm square block of mycelium was cut from the culture near the leading edge and mechanically macerated in an OMNI Bead Ruptor Elite (Onelab, Auckland, NZ) for 30 s at 4 m/s using a 0.5 cm ceramic bead (MP Biomedicals, Auckland, NZ). The macerate (1 mL) was inoculated onto approximately 30 wheat grains in a Petri dish, pre-sterilised according to the method of S. Card [32] and incubated in the dark for 3 weeks at 22°C. Four infected grains (per tube) were stored in cryopreservation tubes containing 500 µL of 30% (v/v) glycerol at-80°C.

### 4.2 Extraction and quantification of sporidesmin A in Pse. chartarum spores

Spores from three technical replicates of *Pse. chartarum* strain 91/35/29, grown on sterile wheat grains as described above, were harvested at 3 and 4 weeks post inoculation to ensure at least one harvest had accumulated spores and sporidesmin A. Sporidesmin was extracted by mixing approximately 50 mg of infected grain in 1 ml of 0.05% (v/v) Tween 20 for 15 min at room temperature on a mini Labroller rotator (Labnet International Inc, NJ, USA). The extract was allowed to settle at room temperature for 5 min, 50 µl of the cleared supernatant was diluted 10 fold in 0.05% (v/v) Tween 20, and sporidesmin A quantified using a direct competitive Enzyme Linked Immunosorbent Assay (ELISA) as described previously [33]. The ELISA has a working range of 0.2- 12.4 ng sporidesmin per ml.

### 4.3. Pse. chartarum visualisation

*Pse. chartarum* spores were cultured on wheat grains for three weeks as already described. A 5 mm segment of an infected grain was embedded in 5% (w/v) water agarose (Difco ™ Becton, Dickinson and Co. USA) and 30 µm sections cut into phosphate-buffered saline (pH 7.4) using a vibratome (Leica VT1000S Vibratome, Wetzlar, Germany). The sections were examined using bright field microscopy on a BX53 microscope fitted with a UPlanfl N 20x/0.5 objective and coupled to a SC180 camera with cellSens standard software (v 2.3) (Olympus, Tokyo, Japan).

### 4.4. Preparation of genomic DNA

*Pse. chartarum* mycelia were grown in four 250 mL flasks containing 50 mL each of potato dextrose broth (Difco™ Becton, Dickinson and Co., NJ, USA) for 5 days at 22°C under a 8 hr light, 16 hr dark cycle. High molecular weight genomic DNA was extracted from the mycelia using the Quick-DNA™ Fungal/Bacterial MidiPrep kit (Zymo Research Corp., CA, USA) according to the manufacturer’s instructions. DNA concentration was quantified using both the Nanodrop ND-1000 spectrophotometer (Thermo Fisher Scientific, MA, USA) and the Qubit fluorometer BR assay (Thermo Fisher Scientific, MA, USA), following the manufacturer’s guidelines.

### 4.5. Genome sequencing and assembly

The *Pse. chartarum* genome was sequenced by Macrogen (Seoul, South Korea) on the RSII platform using single molecule real time (SMRT) long read sequencing technology from Pacific Biosciences (PacBio, CA, USA). The sequencing chemistry included the 20Kb SMRTbell template and the polymerase binding kit P6 Version 2. The instrument produced 450K reads with an average length of 12 Kb, and a total count of 5.550 billion nucleotides. A primary assembly was produced from PacBio RSII reads using CANU Version 1.5 [34] assuming a genome of 35 Mb and with standard parameters. For the purposes of identifying the sporidesmin gene cluster, the completeness of the genome assembly and annotation was estimated by quantifying the expected gene content using the Benchmarking Universal Single-Copy Orthologs (BUSCO) software tool Version 3.0.2 with the “fungi_odb9” database as the reference [35]. To assess bacterial contamination in the *Pse. chartarum* genome, the PacBio reads were aligned to proteins using the “blastx” function of Diamond Version 0.9.14.115 [36], with tabular output but otherwise default settings. The PacBio reads with non-fungal protein alignments were aligned to the genome using Minimap2 Version 2.17-r941 [37], to check genome positioning and coverage. The reads with non-fungal protein alignments all aligned to several shorter (< 10 kb) contigs of the genome and these were removed from the final assembly. It was inferred that there was zero detectable bacterial contamination from the analysis.

### 4.6. Genome annotation and bioinformatics

The genome was annotated using the evidence-based MAKER genome annotation pipeline (Version 2.31.10) [38]. The putative sporidesmin gene cluster was identified using tBLASTn from the BLAST+ suite Version 2.7.1 [41] with a cut-off filter of 1e^-10^. Proteins from the aspirochlorine, sirodesmin and gliotoxin clusters of *A. oryzae, L. maculans* and *A. fumigatus* respectively were used as query sequences as the ETP gene clusters in these species have been functionally characterised [18-20]. The proteins for aspirochlorine and gliotoxin synthesis were obtained from the MIBiG 2.0 database [42], and sirodesmin proteins were obtained from the NCBI protein database. Protein accession numbers are given in Supplementary Tables S2. The domain architecture of inferred proteins from the putative sporidesmin gene cluster was predicted by InterPro Version 7 at http://www.ebi.ac.uk/interpro/ [43] or Pfam Version 32 [44].

### 4.7. Nucleotide sequence accession number

The whole genome sequence of *Pse. chartarum* strain 91/35/29 has been deposited at NCBI’s Sequence Read Archive (SRA) and GenBank under Bioproject number PRJNA596299. The SRA accession number of draft genome of *Pse. chartarum* strain 91/35/29 is PRJNA596299. The putative sporidesmin cluster has been submitted to Genbank under the accession MT353658.

## Acknowledgements

This work was supported by the New Zealand Ministry for Business, Innovation and Employment Strategic Science Investment Fund, through the AgResearch Curiosity Fund [50001]; Genomics Aotearoa [UOOX1702] and the University of Auckland ‘s Agritech Strategic Research Initiatives Fund [9841/3708503].

## Table Legends

**Table S1A**. *E*-values from tBLASTn comparisons of aspirochlorine cluster genes against the *Pse. Chartarum* genome (cut-off is e-^10^).

**Table S1B**. *E*-values from tBLASTn comparisons of sirodesmin cluster genes against the *Pse. Chartarum* genome (cut-off is e-^10^).

**Table S1C**. *E*-values from tBLASTn comparisons of gliotoxin cluster genes against the *Pse. Chartarum* genome (cut-off is e-^10^).

**Table S2:** BLASTp comparisons of sporidesmin toxin cluster genes against the UniprotKB database.

**Table S3**. NCBI accession numbers of LSU sequences and sequenced genomes used in the construction of the phylogenetic tree.

---

**Fig. S1.**
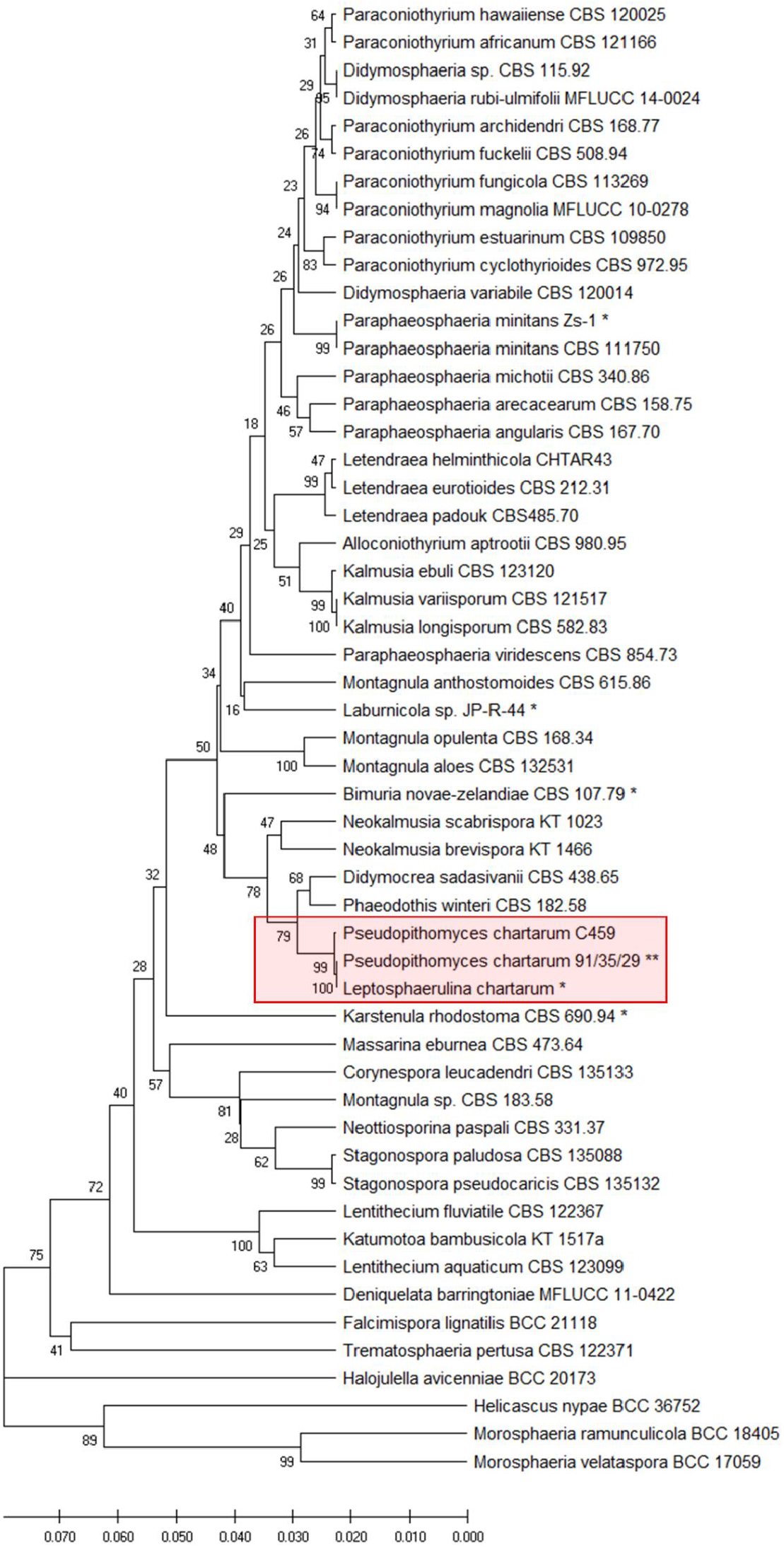
Phylogram of the nucleic acid alignment of the ribosomal LSUs from species of the Pleosporales. The phylogenetic tree was created based on individual LSU sequences from the complete genomes of the Pleosporales listed and recovered from the Genbank (NCBI) database (as of July 2020). The red box highlights the *Pseudopithomyces* clade. A bootstrap of 1,000 replicates was used to establish the percentage support for each branch of the phylogenetic tree [47]. The evolutionary distances were computed using the Maximum Composite Likelihood method [4 8] and the scale bar shows the number of base substitutions per site. An asterisk denotes the strains with a complete genome sequence. The sporidesmin-producing *Pse. chartarum* strain 91/35/29 sequenced in this study is denoted by two asterisks. The LSU sequences without asterisks were originally a ligned as previously described [1], except for *P. chartarum* (C459). The sequences from each gene set was aligned using MAFFT v 7.450 [4 9], by itself and trimmed to keep the core of the alignment, and these were then concatenated into a single fragment for each organism. The accession numbers for individual LSU genes, and the whole genomes from which the others were retrieved, are listed in Supplementary Table S 3.

